# Among-individual variation of risk-taking behaviour in group and solitary context is uncorrelated but independently repeatable in a juvenile Arctic charr (*Salvelinus alpinus*) aquaculture strain

**DOI:** 10.1101/2021.12.06.471387

**Authors:** Joris Philip, Marion Dellinger, David Benhaïm

**Affiliations:** School of Biology, University of St Andrews, St Andrews, U.K.; Department of Aquaculture and Fish Biology, Hólar University, Saudárkrókur, Iceland

**Keywords:** Animal personality, Repeatability, Cross-context consistency, Risk-taking, Individual variation, Welfare, Aquaculture

## Abstract

Behavioural traits have been shown to have implications in fish welfare and growth performances in aquaculture. If several studies have demonstrated the existence of repeatable and heritable behavioural traits (i.e., animal personality), the methodology to assess personality in fishes is often carried out in solitary context, which appears to somewhat limit their use from a selective breeding perspective because these tests are too time consuming. To address this drawback, group-based tests have been developed. In Nordic country, Arctic charr (*Salvelinus alpinus*) is widely used in aquaculture, but no selection effort on behavioural traits has yet been carried out. Specifically, in this study we examined if risk-taking behaviour was repeatable and correlated in group and solitary context and if the early influences of physical environment affect the among-individual variation of behavioural trait across time in order to verify whether a group risk-taking test could be used as a selective breeding tool. Here, we found that in both contexts and treatments, the risk-taking behaviour was repeatable across a short period of 6 days. However, no cross-context consistency was found between group and solitary, which indicates that Arctic charr express different behavioural trait in group and solitary.

## Introduction

Behavioural traits have been shown to have implications in a wide range of biological fields and livestock productions including fish welfare and growth performances in aquaculture (Huntingford et al., 2006). Several studies have demonstrated the existence of repeatable and heritable behavioural traits in the behavioural ecology frameworks i.e., animal personality defined as consistent among-individual variation in average behaviour across repeated measures (Dingemanse et al., 2009; Dochtermann et al., 2019). Consistent behavioural traits have been shown in different fish species such as boldness and aggression in Zebrafish (*Danio renio*; (Ariyomo et al., 2013), boldness in Seabass (*Dicentrarchus labrax*; (Ferrari et al., 2016), boldness and aggression in Brown trout (*Salmo trutta*) (Kortet et al., 2014), suggesting that it may be possible to select individuals on the basis of behavioural traits.

One way to influence the behaviour in a farming context in order to increase welfare is to add complexity to the rearing condition (Huntingford, 2004; Näslund and Johnsson, 2016). The addition of a 3D physical enrichment (i.e., plastic plants and/or stones) can increase the environmental complexity and decrease maladaptive and aberrant traits compared to those observed in fish reared in a plain environment (Macaulay et al., 2021). It is also argued that physical complexity plays a major role in the development of such trait. According to Réale et al (2007) a behavioural trait can be biologically explained by the number of genes involved in the expression of this trait. It simply refers to the fact that the influence of the genetic background in a given environment will affect the expression of a given trait (i.e., Risk-taking). Indeed, the environmental influences at an early stage of development could affect the expression of a behavioural traits later in the lifespan (Stamps and Groothuis, 2010). Nevertheless, the effect of gene and environment could not be seen independently when the genotype of individuals is divergent (i.e., different populations in different environments) because both affect the expression of personality (i.e., among-individual consistency in behavioural trait) (Réale et al., 2007; Stamps and Groothuis, 2010). Domesticated species tend to share the same genotype across selected generations but could however experience different environmental conditions and this is precisely where they are likely to express different behavioural traits (Cabrera et al., 2021; Castanheira et al., 2017; Ferrari et al., 2016; Johnsson et al., 2014; Stamps and Groothuis, 2010).

Several methodological approaches have been used to assess personality in fishes, including individual-based tests such as confinement in rainbow trout (*Oncorhynchus mykiss*; (Øverli et al., 2007, 2004), feeding recovery in a novel environment in African catfish (*Clarias gariepinus*; (Martins et al., 2006)) and the Nile tilapia (*Oreochromis niloticus*; (Martins et al., 2011)), exposure to a novel object in Nile tilapia (Martins et al., 2011), aggression tests in rainbow trout (*Onchorynchus mykiss*; (Øverli et al., 2007)) or boldness in Atlantic salmon (*Salmo salar*; (Benhaïm et al., 2020)). Most behavioural tests are therefore carried out in isolation conditions which appears to somewhat limit their use from a selective breeding perspective because these tests are too time consuming. To address this drawback, group-based tests have been developed. Most of these tests concern risk-taking in Seabass (*Dicentrarchus labrax*; (Ferrari et al., 2015)) or common carp (*Cyprinus carpio*; (Huntingford et al., 2010)). However, the link between individual- and group-based tests has not yet been clearly established. No link has for example been found in Seabass and Gilt head seabream (Castanheira et al., 2013; Ferrari et al., 2015; Millot et al., 2009).

It appears essential to characterize risk-taking behaviour in order to evaluate the potential ability of the targeted fish species to cope with potentially stressful conditions in their rearing conditions and to ensure good welfare conditions. Furthermore, selective breeding is widely practiced in fishes and selection has often been applied to growth as a major trait of interest. However, the distribution of behavioural traits within a group may play an important role in the welfare of reared fish (Adams and Huntingford, 2005). Developmental aspects such as raising conditions are of fundamental interest for behavioural traits and there are potential links between risk-taking and growth performances. Indeed, risk-taking as well as other behavioural traits are known to influence growth parameters (Biro and Stamps, 2008). A link has been found between self – feeding behaviour and coping - styles (in Tilapia and Seabass (Benhaïm et al., 2017; Ferrari et ùal., 2014)), as well as a link between individual growth performance and various behavioural traits (Ferrari et al., 2016; Huntingford et al., 2010; Sundström et al., 2004). The link between individual performances and behavioural traits has been found to be context and/or species dependent. For example, boldness and swimming activity were positively correlated with growth rate in common Sole (*Solea solea*; (Mas-Muñoz et al., 2011), whereas a negative correlation was found in Seabass, where shyer fish exhibited higher growth rate in a predictable environment in terms of food supplies. In studies on common carp, seabass and seabream, metabolic rate was found to be significantly higher in risk – taking fish (Herrera et al., 2014; Huntingford et al., 2010; Jenjan et al., 2013; Killen et al., 2011) whereas in species with a passive benthic lifestyle, such as the Senegalese sole (*Solea senegalensis*), bold individuals were shown to consume less oxygen (Martins et al., 2011). Therefore, there is clearly a need to develop tools that assess the behavioural among-individual variation in farmed species to improve their welfare and likely their farmed value.

In Nordic countries, Arctic charr (*Salvelinus alpinus*) is widely used in aquaculture for food production and represents a considerable economic value (Imsland et al., 2019). This production is the result of selective breeding programs run over the last 30 years with a focus on growth rate, feed conversion, survival rate, size, and age at maturation (Olk et al., 2019). However, no selection effort on behavioural traits has yet been carried out. In the present work, the effect of physical enrichment on the among-individual variation response in risk-taking behaviour was tested. Specifically, we compared this behavioural trait in an individual and a group-test over a short-term period in order to compare the among-individual variation in risk-taking consistency in both contexts and to verify whether there are correlated to each other. We predicted (1) risk-taking behaviour to be repeatable in individual and group-based tests, (2) the existence of cross-context consistency in Arctic charr i.e., correlation between individual and group-based tests and, the influence of 3D physical environment on risk-taking behaviour repeatability in both contexts.

## Material & methods

### Biological model and housing

An Icelandic aquaculture strain of Arctic charr from the Hólar University breeding program (Iceland) was used in this experiment. Eggs were incubated at 4°C in a flow through system. After hatching, the batch was split into six 20L-tanks with a biomass of 13.8 g per tank (i.e., 644 g.m^3^), which gave three replicates of each treatment. After first feeding (73 days post hatching; dph) three of the tanks were enriched in three dimensions for the environmental enrichment treatment: i.e., vertically with a green plastic plant, and horizontally with five black volcanic rocks. The temperature in the tanks was maintained at 5 ± 1 °C over the course of the experiment. The fish were fed three times a day (9:00, 13:00, 16:00) with commercial aquaculture pellets according to the Inicio guidelines (BioMar). The waterflow was 48 L.hour^-1^ at hatching and increased gradually to 120 L.hour^-1^ to keep the oxygen level at 100% of saturation. The batch was on a 12:12-hour light schedule.

### Sampling and tagging

A randomly selected sample of 32 individuals per tank was tagged. The tagging was done under anaesthesia (2-phenoxyethanol) at a concentration of 310 ppm at 259 dph. A small incision was made between the pectoral fins and a PIT-tag (Passive Integrated Transponder 1.4 x 8mm FDX-B PIT tags, Oregon RFID EU GmbH) was inserted into the peritoneal cavity. Behavioural tests started 170 days after tagging, hence fish had totally recovered from the surgery at that time.

### Experimental design

Solitary and group risk-taking were each assessed twice, in an OFTS (i.e., Open Field Test with Shelter) and an GRT (i.e., Group Risk taking Test), respectively. The OFTS 1 (i.e., first trial repetition – R1) was assessed between 429 and 435 dph and the OFTS 2 (i.e., R2) between 436 and 452 dph. The GRT 1 was assessed between 453 and 459 dph and GRT 2 between 453 and 465 dph.

#### Risk-taking

##### Solitary risk-taking

To assess the solitary risk-taking (SRT), an OFTS was used. The tests were carried out in a rectangular arena (29.5 cm x 39.7 cm) made of opaque white Plexiglas® with a shelter (6 cm x 14 cm) in its left corner (Figure 1). This non-forced test allowed individuals to stay hidden in the shelter. The decision to exit the shelter into the open area (i.e., exploration zone) was controlled by the individuals. The latency time to emerge from the shelter (s) was measured as a proxy of risk-taking. The OFTS arena was situated above a white LED backlight (110 x 110 cm, Noldus, the Netherlands). A camera (Basler Ace acA1920-150 mm camera Germany, 30 fps) was located 112 cm above the arenas and plugged into a computer. The videos were recorded using the Ethovision XT15 tracking software (Noldus, The Nertherlands. The selected individuals were placed into the shelter through a roof door. After 5 minutes of acclimation the front door of the shelter was lifted, and the individuals were free to move around in the arena for 20 minutes. After 20 minutes the individuals were caught and anesthetized with 2-phenoxyethanol at a concentration of 310 ppm. The weight and the fork length were recorded, and the individuals were returned to their home tank after full anaesthesia-recovery in clear freshwater.

##### Group risk-taking test

To assess the among-individual risk taking in group, a Group Risk taking Test (GRT) adapted from Ferrari et al (2016) was used. The test was carried out in a 0.114 m^3^ rectangular tank divided into three rooms (Figure 2): a dark shelter-room (0.0314 m^3^) covered by a black lid, a passing tunnel (0.0022 m^3^) and a risk-room (0.0314 m^3^). The device was in a flow through system (100 L.h^-1^). A white LED light was situated above the passing-tunnel and the risk-room. A PIT-tag reader circular antenna (diameter: 10 cm, Dorset) linked with an USB data logger (Dorset) was placed in the middle and around the tunnel. A group of 32 individuals were placed into the shelter-room by the roof door and acclimatized for 60 min. At the end of the acclimation time, the shelter-room front door was gently lifted, and the individuals were free to pass through the tunnel to access the risk-room for 24 hours (1440 min). The latency to first exit (s) was recorded. When the individual swam through the antenna, the PIT-tag number of the fish, the date and the time was recorded by the USB data logger. After the test, the individuals were caught and anesthetized with 2-phenoxyethanol at a concentration of 310 ppm. The weight and the fork length were recorded, and the individuals returned to their home tank after full anaesthesia-recovery in clear freshwater.

**Figure 2:**
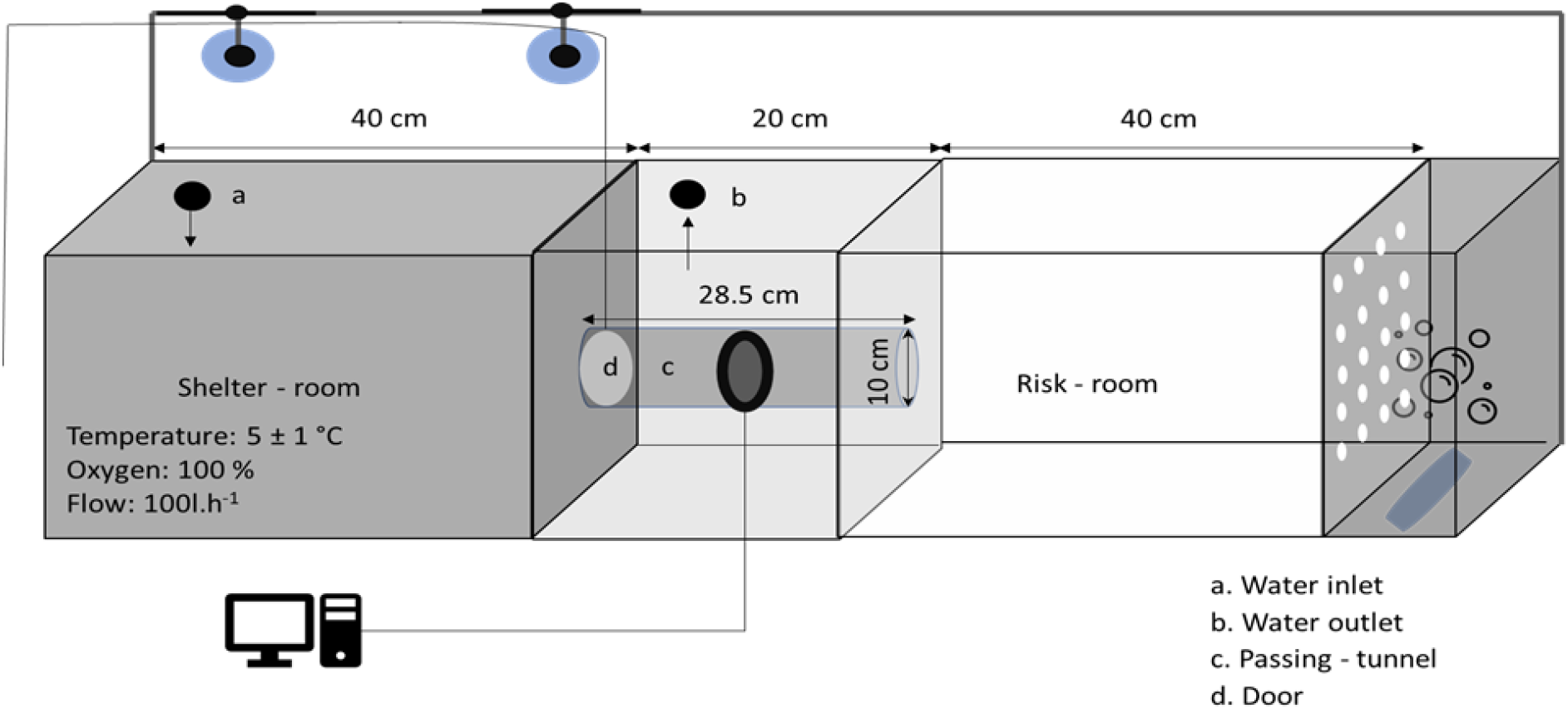
Group risk-taking device. The water inlet (a) is situated in the shelter room and the outlet (b) in the passing room. To prevent from unbalanced oxygen concentration throughout the rooms, an oxygenation was constantly running in a back of the risk – room. The door was tied to a rope through two pulleys. When the acclimation time was over, the door was lift through the rope.

#### Statistical analysis

All statistical analyses were performed using R v.1.4.1103 R Core Team, 2020. The assumptions for a linear mixed model and linear model (i.e., normality of residuals, homoscedasticity of the residuals and uncorrelated residuals) were examined and validated for each model.

### Repeatability of risk-taking

The repeatability was assessed using the package and function rptR (Nakagawa and Schielzeth, 2010; Stoffel et al., 2017). This function allowed the use of a linear mixed model (LMM) and extract the variances of interest in order to calculate the repeatability estimates using Equation 1. When the repeatability is calculated using the function, it is reasonable to follow a normal distribution for the response variable (Dochtermann and Dingemanse, 2013). A logarithm transformation of the latency in SRT and GRT was applied to fit a gaussian distribution. In the model, the logarithm of the latency to first exit of the shelter was the response variable, and the trial repetition was a fixed factor. The body mass was used as a covariable. The individual ID was used as a random factor. The parameters nboot and npermut allowed the formula to calculate the confidence intervals by random iterations and were set at 1000.

### Among-individual correlation of risk-taking in solitary- and group-based test

To assess the among-individual correlation of risk-taking behaviour between solitary- and group-based tests, generalized linear mixed model using Bayesian Markov Chain Monte Carlo (MCMC) methods were used with a non-informative parameter-expanded prior (Hadfield et al., 2010; Houslay and Wilson, n.d.). A bivariate model was used to associate SRT with GRT (i.e., joint response variables). The fixed effects were the trial repetition and the body mass. The individual ID was used as a random factor and defined as an unstructured covariance matrix in order to calculate the among-individual variance for the two response variables separately and for the covariance between them. Number of iterations were 420000; burn-in=20000; thin=100. To calculate the among-individual correlation (r) between the two risk taking tests, the posterior distribution of (co)variance between traits (i.e., cov(SRT, GRT)) were divided by the product of the square root of their variances (Equation 2). The mean correlation estimate was extracted with the function mean and the confidence intervals with the function HPDinterval. Here, as the correlation estimates can take on either positive or negative values, the credible intervals of correlation of the (co)variance (i.e., with standardized covariance to a scale of −1 to 1) were used to assess statistical significance (95% credible interval > 0).

## Results

### Repeatability of risk-taking

The repeatability estimates showed a significant consistency in both treatments and both risk-taking behaviours at short-time scale (Table 1; Enriched: SRT, R= 0.51 ± 0.084 [0.32, 0.68], GRT, R = 0.50 ± 0.092 [0.36, 0.63]. Plain: SRT, R= 0.50 ± 0.095 [0.43, 0.72], GRT, R = 0.45 ± 0.117 [0.24, 0.59]).

**Table 1:** Correlation estimates (r) and confidence intervals (CI) of group and risk-taking behaviours between treatments (i.e., enriched, and plain).

| **Trait** | **Treatment** | **r** | **CI** |
| --- | --- | --- | --- |
| SGR, GRT | Enriched | -0.22 | [-0.56, 0.17] |
|  | Plain | 0.15 | [-0.29, 0.63] |

### Risk – taking correlation

No association was found between SRT and GRT in either treatment. The correlation estimates were very low, and the confidence intervals spread between −1 and +1 (Enriched r= −0.22 [−0.56, 0.17], Plain: r= 0.15 [−0.29, 0.63]) (Figure 3, Table 2).

**Figure 3:**
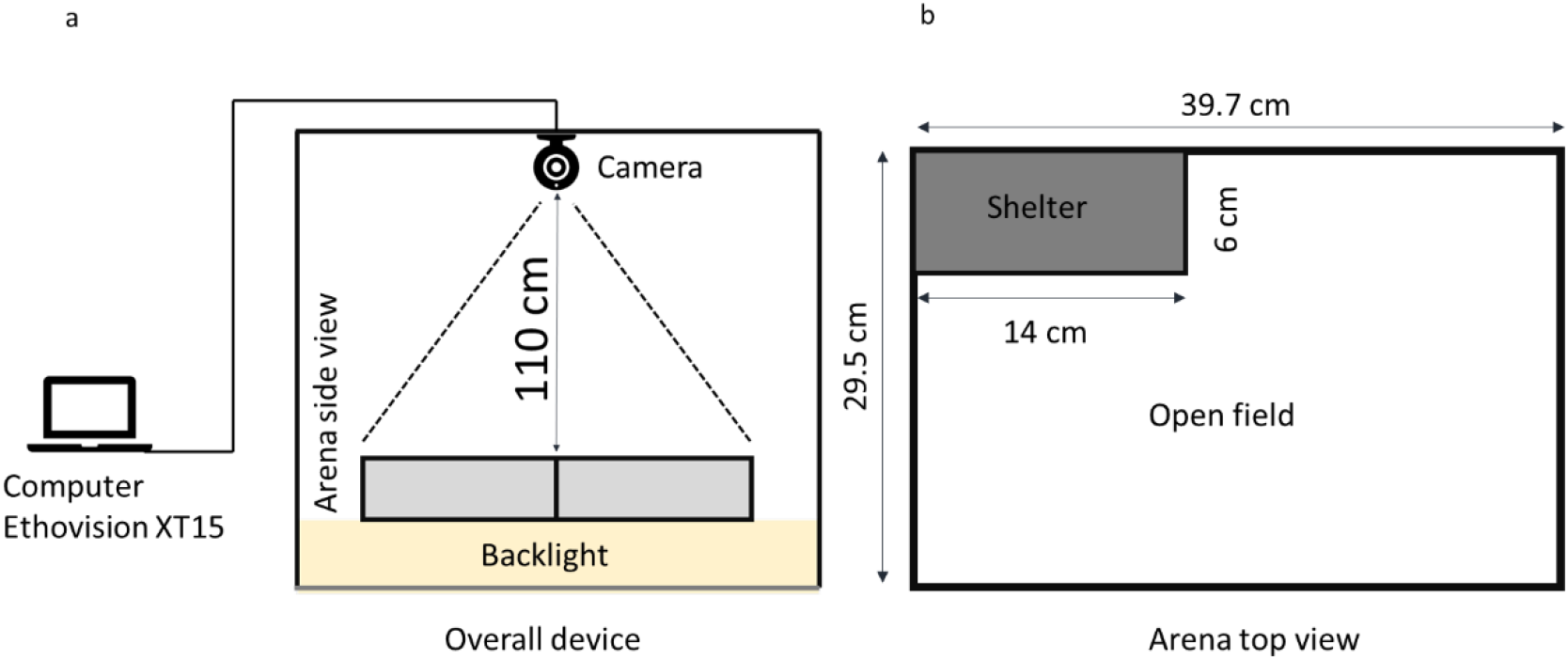
Open Field Test with Shelter device. a. Front view of the overall device. b. Schematic (top view) of the arena.

**Figure 4:**
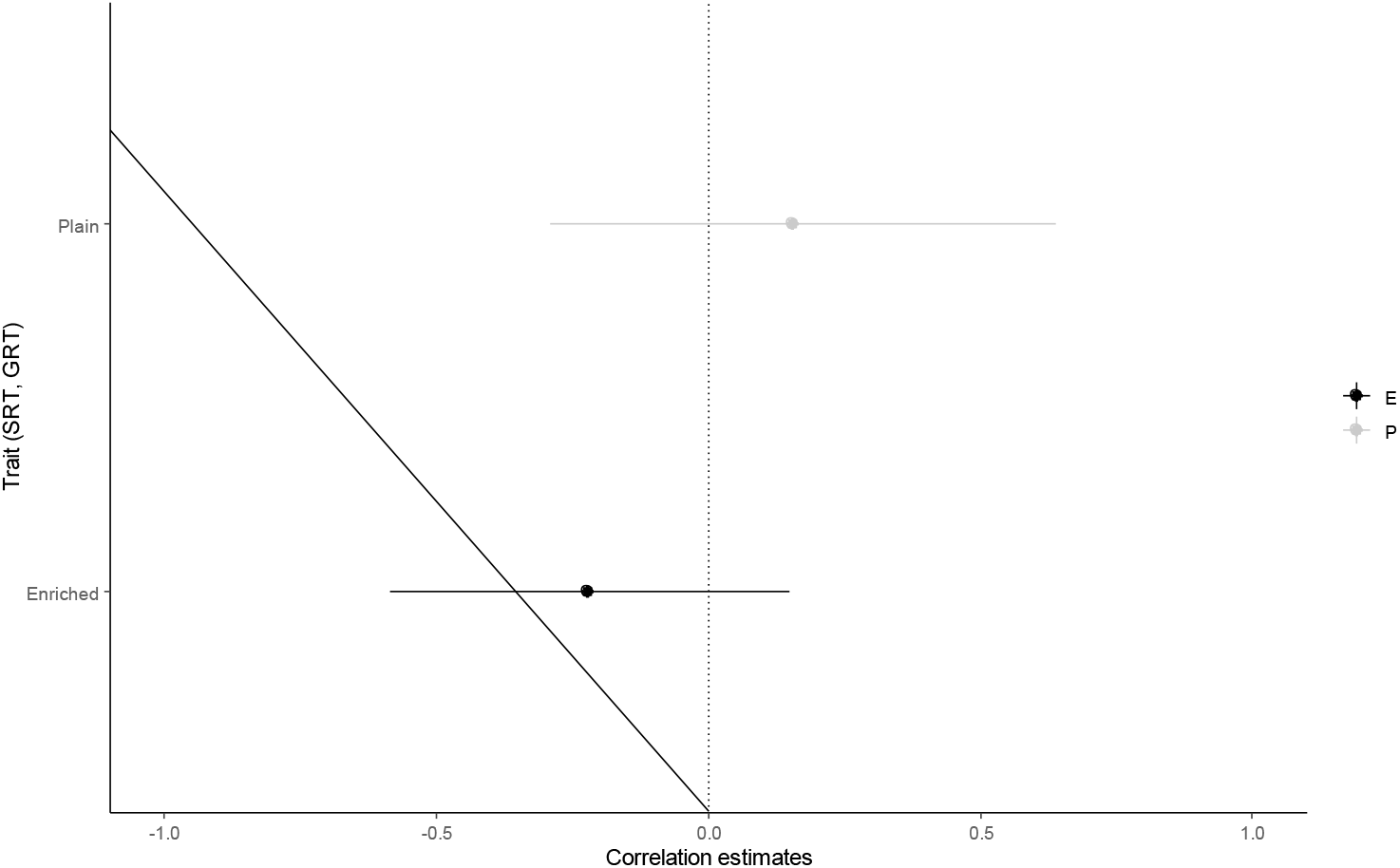
Correlation estimates of trait association. The point is the mean estimate and the line the confidence interval.

**Table 2:** Repeatability estimates (R), standard error (SE) and confidence intervals (CI) in both risk-taking behaviours (i.e., solitary and group risk-taking, respectively SRT and GRT) and treatments (i.e., enriched, and plain).

| **Trait** | **Treatment** | **R** | **SE** | **CI** |
| --- | --- | --- | --- | --- |
| SRT | Enriched | 0.51 | 0.084 | [0.329, 0.683] |
|  | Plain | 0.50 | 0.095 | [0.433, 0.723] |
| GRT | Enriched | 0.50 | 0.092 | [0.362, 0.639] |
|  | Plain | 0.45 | 0.117 | [0.243, 0.592] |

## Discussion

The aim of this work was to assess the effect of two different contexts (i.e., solitary, and group) and treatments (i.e., enriched, and plain) on risk-taking behaviour. We examined if risk-taking behaviour was repeatable and correlated in different contexts across time in order to verify whether a GRT could be used as a selective breeding tool. In both contexts and treatments, the risk-taking behaviours were repeatable across a short period of 6 days. However, no correlation was found between the two contexts, suggesting that there are differences in risk-taking behaviours depending on the setting.

The results of this work are in accordance with those of previous studies that also showed consistency across a short-term period in individual context e.g., Seabass (Ferrari et al., 2014; Millot et al., 2009), Common carp (Huntingford et al., 2010) as well as in group-based context e,g, Seabass (Ferrari et al., 2015) and Gilthead seabream (Castanheira et al., 2013). It is therefore likely that risk-taking is repeatable across time in both contexts. Nevertheless, risk-taking was not cross-context consistent which indicates that different behavioural traits were expressed in each situation. An explanation for this difference is the influence of social interactions that are occurring when risk–taking behaviour is measured in a group context, which has been found in other studies. For example, no cross-context consistency has been found in Seabass despite independent repeatability in each context (Castanheira et al., 2013; Ferrari et al., 2015; Millot et al., 2009). The fact that cross–context correlations among behavioural traits is a fundamental aspect of animal personality theory, cross-context consistency in a behavioural trait is rarely found between individual and group-based contexts. In fact, cross-context consistency between individual and group-based tests has only been found in Gilthead seabream where group response to hypoxia was found to be correlated with individual feeding recovery (Castanheira et al., 2013).

Social interactions can be extremely complex in fish species and particularly in salmonids. It is likely that the relationships between individuals are dependent on the behavioural rank of each individual within a group. In other words, group organisation is mediated by among-individual variation in behaviour (Croft et al., n.d.); for example, bolder individuals become more dominant and shyer individuals become subordinates. Indeed, it is known that Arctic charr is a very territorial species with often one or two dominant individuals in one group (Adams et al., 1995). Therefore, one scenario that could explain the lack of cross-context consistency between individual and group contexts is the strong influence of social interactions in Arctic charr and a need for subordinates to escape more dominant individuals in a group setting.

In this experimental setup, the group was at low density, and it is possible that an individual exiting the shelter-room does not express a risk-taking behaviour but rather escapes to the safe area to avoid dominant chase and aggression in the shelter-room i.e., the shy fish exit the safe area before the bold ones. Although these results do not explicitly support this hypothesis because no negative correlations between the individual and group were found, it seems likely that aggression could have been a factor influencing the exit behaviour (personal observation). Therefore, the time to exit the safe area could be somehow linked to the fish social rank, and it would be interesting to follow this idea further. This idea is also supported by studies on other salmonid species where the social rank could be correlated with risk-taking, i.e., risk-taking-aggressiveness syndrome as in Brown trout (Sundström et al., 2004) and Atlantic salmon (Adams and Huntingford, 2005).

Nevertheless, other scenarios where social learning and leadership could have driven exit behaviour could be put forward. These two theories are complementary to each other as social leaning refers to learning influenced by observation of other congeners, and this could lead to leadership where the initiation of a movement, made by one or some individuals is followed by the rest of the group (Galef and Giraldeau, 2001; Krause et al., 2000). Therefore, small groups of sociable (i.e., unrelated to boldness) individuals could have influenced the exit behaviour.

In conclusion, in the present study individual and group risk-taking behaviour were independently repeatable. However, no cross-context consistency was found, therefore, the group risk-taking test as designed in this experiment cannot be used to select for boldness in the Arctic charr. We also observed that the rearing condition did not have any effect on the among-individual consistency or on the cross-context consistency. The theory of the development of among-individual consistency suggests that a series of continuous interactions between internal factors (i.e., genetic, and neural activity) and contextual factors (i.e., environment) induces among – individual behavioural consistency (i.e., animal personality). Personality trait expression can be the result of epigenetic modifications (Stamps and Groothuis, 2010) based on a set of molecular processes altering genes activity and the maintenance of genetic activity variation mostly occurs through genotype by environment interaction (Réale et al., 2007; Stamps and Groothuis, 2010). Here, the fact that we did not find any difference in the among-individual structure of risk-taking between the two treatments suggest that genetic factors explained most of the behavioural phenotype and/or that the enrichment did not induce any developmental change over the period of this experiment. Further research is needed to better understand the link between individual and group behaviour. Indeed, it seems that social interaction strongly affects the risk-taking behaviour resulting in a biased behavioural measure (i.e., exit behaviour is not always a reliable risk-taking behaviour proxy). It is important to note that the measured behaviour is independently repeatable in both tests, which indicates that Arctic charr express different behavioural trait in group and solitary context. We therefore recommend further investigation into group and solitary risk-taking and social interaction (i.e., aggression, social learning, and leadership) in order to disentangle the real among-individual variation of risk-taking behaviour in the Arctic charr.

## Acknowledgements

The authors are very grateful to Pr. Colin Adams and Dr. Marie Laure Bégout for critical comments and advises regarding the conception of the study as well as the original draft of this paper. We would like to thank Marloes Bon for her small participation to this study as part of her BSc project as well as Léo Suret for his precious help during the behavioural assessment. We wish to address many thanks to the staff of the Holar University and specially to Kari Einar and Amber Monroe for taking care of the fish husbandry. This work was funding by Rannis.

## Credit authorship contribution statement

**Joris Philip:** Investigation, GRT experimental device, Data curation, Statistical analysis, Writing original draft. **Marion Dellinger:** Data curation, Review-original draft. **David Benhaïm:** Funding acquisition, Supervision, Data curation, Review and editing.

## Equations

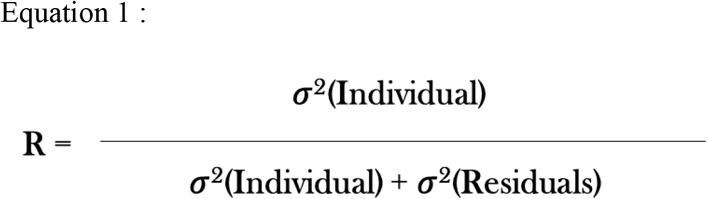

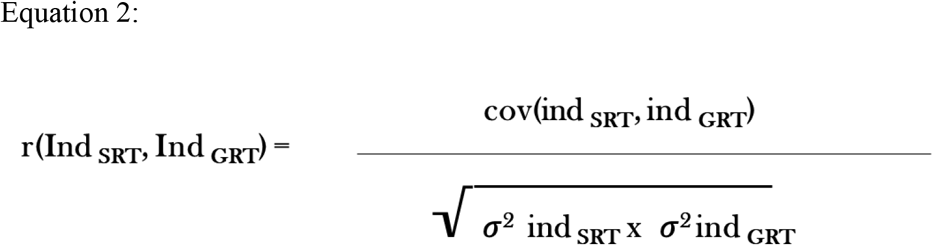

